# Aubergine and piRNAs repress the proto-oncogene *Cbl* for germline stem cell self-renewal

**DOI:** 10.1101/134304

**Authors:** Patricia Rojas-Ríos, Aymeric Chartier, Martine Simonelig

**Affiliations:** mRNA Regulation and Development, Institute of Human Genetics, UMR9002 CNRS-Université de Montpellier, 141 rue de la Cardonille, 34396 Montpellier Cedex 5, France

**Keywords:** CCR4-NOT deadenylation complex, *Cbl* proto-oncogene, germline stem cells, piRNAs, PIWI proteins, translational control

## Abstract

PIWI proteins have essential roles in germ cells and stem cell lineages. In *Drosophila*, Piwi is required in somatic niche cells and germline stem cells (GSCs) for GSC self-renewal and differentiation. Whether and how other PIWI proteins are involved in GSC biology remains unknown. Here, we show that Aubergine (Aub), another PIWI protein, is intrinsically required in GSCs for their self-renewal and differentiation. Aub loading with piRNAs is essential for these functions. The major role of Aub is in self-renewal and depends on mRNA regulation. We identify the *Cbl* proto-oncogene, a regulator of mammalian hematopoietic stem cells, as a novel GSC differentiation factor. Aub represses *Cbl* mRNA translation for GSC self-renewal, and does so through recruitment of the CCR4-NOT complex. This study reveals the role of piRNAs and PIWI proteins in translational repression for stem cell homeostasis and highlights piRNAs as major post-transcriptional regulators in key developmental decisions.

## Introduction

The regulation of gene expression at the mRNA level is fundamental for many biological and developmental processes. In recent years, Piwi-interacting RNAs (piRNAs) have emerged as novel key players in the regulation of gene expression at the mRNA level in several models. These 23-to 30-nucleotide-long non-coding RNAs are loaded into specific Argonaute proteins, the PIWI proteins (Guzzardo et al, 2013; Ishizu et al, 2012). Classically, piRNAs repress transposable element expression and transposition in the germline. They are largely produced from transposable element sequences and target transposable element mRNAs by complementarity, which induces their cleavage through the endonuclease activity of PIWI proteins and represses their expression.

Recent studies have shown that piRNAs also target protein-coding mRNAs, leading to their repression by PIWI-dependent mRNA cleavage or via the recruitment of the CCR4-NOT deadenylation complex. This regulation is required for embryonic patterning in *Drosophila* (Barckmann et al, 2015; Rouget et al, 2010), sex determination in *Bombyx mori* (Kiuchi et al, 2014), and degradation of spermiogenic mRNAs in mouse sperm (Goh et al, 2015; Gou et al, 2014; Watanabe et al, 2015; Zhang et al, 2015).

piRNAs involved in the regulation of protein-coding mRNAs in *Drosophila* embryos are produced in the female germline and provided maternally. An open question is whether this function of the piRNA pathway in post-transcriptional control of gene expression plays a role in the biology of germ cells and germline stem cells (GSCs). In the *Drosophila* ovary, two to three GSCs are localized in the anterior-most region of each ovariole and self-renew throughout the adult life, giving rise to all germ cells. GSCs in contact with somatic niche cells divide asymmetrically to produce a new stem cell that remains in contact with niche cells (self-renewal) and another cell that differentiates into a cystoblast, upon losing the contact with the niche. Subsequently, the cystoblast undergoes four rounds of synchronous division with incomplete cytokinesis to produce a cyst of sixteen interconnected germ cells, of which one cell is specified as the oocyte and the other fifteen cells become nurse cells (Figure 1A).

**Figure 1:**
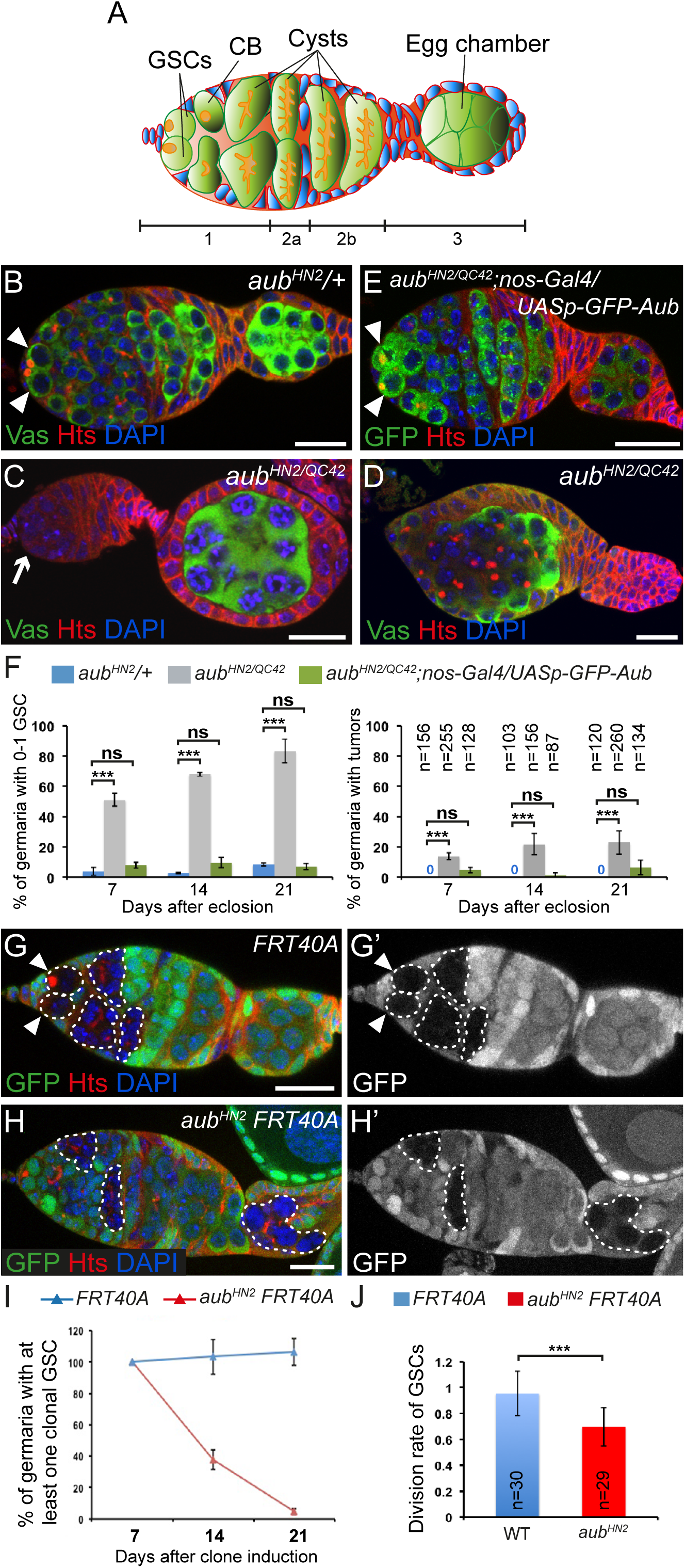
Intrinsic role of Aub in GSC self-renewal and differentiation. (A) Schematic diagram of a germarium showing the somatic cells (blue) and the germline cells (green). The spectrosomes and fusomes are shown in orange. The different regions of the germarium are indicated. Region 1: dividing cysts; region 2: selection of the oocyte; region 3: egg chamber with posteriorly localized oocyte. GSCs: germline stem cells; CB: cystoblast. (B-E) Immunostaining of germaria from 7 day-old females with anti-Vasa (green) and anti-Hts (red). DAPI (blue) was used to visualize DNA. (B) *aub*^*HN2*^/+ was used as a control. (C, D) examples of *aub*^*HN2/QC42*^ germ cell loss and tumor, respectively. (E) Phenotypic rescue of *aub*^*HN2/QC42*^ with *UASp-GFP-Aub* expressed using *nos-Gal4*. White arrowheads indicate GSCs; the white arrow indicates GSC loss. (F) Quantification of mutant germaria with 0 to 1 GSC, or with GSC tumors shown in (B-E). The number of scored germaria (n) is indicated on the right graph. *** *p*-value <0.001, ns: non significant, using the χ2 test. (G-H’) Germaria containing control (G, G’) or *aub*^*HN2*^ mutant (H, H’) clonal GSCs stained with anti-GFP (green) and anti-Hts (red), 14 days after clone induction. DAPI (blue) was used to stain DNA. Clonal cells are marked by the lack of GFP. Clonal GSCs and cysts are outlined with dashed line. White arrowheads show clonal GSCs in the control. *aub* mutant clonal GSCs have been lost (H, H’). Scale bar: 10 μm for B-E and G-H’. (I) Quantification of germaria containing at least one clonal GSC at 7, 14 and 21 days after clonal induction. (J) Division rate of wild-type and *aub*^*HN2*^ clonal GSCs. The number of scored germaria (n) is indicated. *** *p*-value <0.001 using the Student’s *t*-test.

Two features make Piwi unique with respect to the other two *Drosophila* PIWI proteins, Aubergine (Aub) and Argonaute 3 (Ago3). First, it represses transposable elements at the transcription level through a nuclear function, whereas Aub and Ago3 act by endonucleolytic cleavage of transposable element mRNAs in the cytoplasm; and second, it plays a role in the somatic and germ cells of the ovary, whereas *aub* and *ago3* function is restricted to germ cells. *piwi* function in GSC biology has long been addressed. *piwi* is required in somatic escort cells (which surround GSCs) for GSC differentiation, as well as intrinsically in GSCs for their maintenance and differentiation (Cox et al, 1998; Cox et al, 2000; Jin et al, 2013; Ma et al, 2014). One molecular mechanism underlying Piwi function in GSC biology has recently been proposed to involve its direct interaction with Polycomb group proteins of the PRC2 complex, leading to indirect massive gene deregulation through reduced PRC2 binding to chromatin (Peng et al, 2016). Regulation of *c-Fos* by Piwi at the mRNA level in somatic niche cells has also been reported to contribute to the role of Piwi in GSC maintenance and differentiation (Klein et al, 2016).

Translational control acting intrinsically in GSCs plays a major role in the switch between self-renewal and differentiation. Two molecular pathways ensure GSC self-renewal through translational repression of differentiation factor mRNAs: the microRNA pathway, and the translational repressors Nanos (Nos) and Pumilio (Pum) (Slaidina & Lehmann, 2014). Nos and Pum bind to and repress the translation of mRNAs that encode the differentiation factors Brain tumor (Brat) and Mei-P26, through the recruitment of the CCR4-NOT deadenylation complex (Harris et al, 2011; Joly et al, 2013). In turn, cystoblast differentiation depends on Bag of marbles (Bam), the major differentiation factor forming a complex with Mei-P26, Sex lethal (Sxl) and Benign gonial cell neoplasm (Bgcn) to repress *nos* mRNA translation; Pum interacts with Brat in these cells to repress the translation of mRNAs encoding self-renewal factors (Harris et al, 2011; Li et al, 2012; Li et al, 2009b; Li et al, 2013).

Aub has a distinctive role in protein-coding mRNA regulation. In the early embryo, Aub binds several hundred maternal mRNAs in a piRNA-dependent manner and induces the decay of a large number of them during the maternal-to-zygotic transition (Barckmann et al, 2015). Aub-dependent unstable mRNAs are degraded in the somatic part of the embryo and stabilized in the germ plasm. These mRNAs encode germ cell determinants, indicating an important function of Aub in embryonic patterning and germ cell development. Indeed, Aub recruits the CCR4 deadenylase to *nos* mRNA and contributes to its deadenylation and translational repression in the somatic part of the embryo. This Aub-dependent repression of *nos* mRNA is involved in embryonic patterning (Rouget et al, 2010).

Here, we address the role of *aub* in GSC biology. We show that *aub* is autonomously required in GSCs for their self-renewal. This *aub* function is independent of *bam* transcriptional repression in the GSCs and activation of the Chk2-dependent DNA damage checkpoint. Using an Aub point-mutant form that cannot load piRNAs, we show that piRNAs are required for GSC self-renewal. Genetic and physical interactions indicate that Aub function in GSCs involves interaction with the CCR4-NOT deadenylation complex. Importantly, we identify *Casitas B-cell lymphoma* (*Cbl*) mRNA as a target of Aub in GSCs. *Cbl* acts either as a tumor suppressor or a proto-oncogene depending on its mutations, which lead to myeloid malignancies in humans (Sanada et al, 2009). *Cbl* encodes an E3 ubiquitin ligase that negatively regulates signal transduction of tyrosine kinases; it plays a role in hematopoietic stem cell homeostasis, maintaining quiescence, and preventing exhaustion of the stem cell pool (An et al, 2015). We show that Aub acts to maintain a low level of Cbl protein in GSCs, and that this repression of *Cbl* mRNA by Aub is essential for GSC self-renewal. Furthermore, we find that *Cbl* is required for GSC differentiation, thereby identifying a role for Cbl in the regulation of yet another stem cell lineage.

This study reveals the function of Aub and piRNAs in GSC self-renewal through the translational repression of *Cbl* mRNA, thus highlighting the role of the piRNA pathway as a major post-transcriptional regulator of gene expression in key developmental decisions.

## Results

### *aub* is intrinsically required in GSCs for their self-renewal and differentiation

Although Aub and Ago3 are expressed in GSCs, their function in GSC biology has not yet been addressed (Brennecke et al, 2007; Gunawardane et al, 2007). GSCs can be recognized by their anterior localization in the germarium as well as the presence of the spectrosome, an anteriorly localized spherical organelle in contact with the niche, which is enriched in cytoskeletal proteins (Figure 1A). Cystoblasts also contain a spectrosome that is randomly located in the cell, whereas cells in differentiating cysts are connected by the fusome, a branched structure derived from the spectrosome (Figure 1A). GSCs and differentiating cells were analyzed by immunostaining with an anti-Hts antibody that labels the spectrosome and fusome, and anti-Vasa, a marker of germ cells.

We used *aub*^*HN2*^ and *aub*^*QC42*^ strong or null alleles (Schupbach & Wieschaus, 1991) to address the role of Aub in GSC biology. Immunostaining of ovaries with anti-Hts and anti-Vasa revealed strong defects in both GSC self-renewal and differentiation, in *aub*^*HN2/QC42*^ mutant ovaries at 7, 14 and 21 days. A large proportion of *aub*^*HN2/QC42*^ germaria had 0 to 1 GSC indicating GSC loss, and this defect increased over time (Figure 1B, C, F). A lower proportion of germaria showed differentiation defects, observed as tumors containing undifferentiated cells with spectrosomes (Figure 1D). This phenotype did not markedly increase with time (Figure 1F). Both *aub*^*HN2/QC42*^ phenotypes were almost completely rescued following expression of GFP-Aub with the germline driver *nos-Gal4*, indicating that both defects were due to *aub* loss of function in germ cells (Figure 1E, F).

Because GSC loss was the most prominent defect in *aub* mutant ovaries, and this defect was not rescued by a Chk2 kinase mutant (see below), we focused on this phenotype. We used clonal analysis as an independent evidence to confirm the intrinsic role of *aub* in GSCs for their self-renewal. Wild-type and *aub* mutant clonal GSCs were generated using the FLP-mediated FRT recombination system (Golic & Lindquist, 1989) and quantified at three time points after clone induction. We first verified that *aub* clonal GSCs did not express Aub (Supplementary Figure 1A, A’). The percentage of germaria with wild-type clonal GSCs was stable over time (Figure 1G, I). In contrast, the percentage of germaria with *aub* mutant clonal GSCs strongly decreased with increasing time after clone induction, showing that *aub* mutant GSCs cannot self-renew (Figure 1H-I). The presence of *aub* mutant clonal differentiated cysts marked with fusomes indicated that *aub* mutant GSCs were lost by differentiation (Figure 1H). To confirm this conclusion, we used anti-cleaved Caspase 3 staining to record cell death and address whether the loss of *aub* mutant GSCs could be due to apoptosis. The number of GSCs expressing cleaved Caspase 3 was low and similar in control (*aub*^*HN2*^*/+*) and *aub* mutant GSCs (Supplementary Figure 1B-D), indicating that *aub* mutant GSCs did not undergo cell death.

Next, we asked whether the GSC self-renewal defect in *aub* mutant ovaries could result from Bam expression in GSCs. Anti-Bam immunostaining of *aub* mutant ovaries demonstrated that *aub* mutant GSCs did not express Bam (100% of n=90, Supplementary Figure 1E-F’).

Finally, we determined the division rate of *aub*^*HN2*^ GSCs by counting the number of cysts produced by a clonal marked mutant GSC and dividing it by the number of cysts produced by a control unmarked GSC in the same germarium (Jin & Xie, 2007). As expected, the division rate of wild-type GSCs (FRT40A chromosome) was close to 1 (0.95), whereas that of *aub*^*HN2*^ mutant GSCs was 0.67, indicating a slower division rate in *aub* mutant GSCs (Figure 1J).

To address the role of Ago3 in GSC biology, we used the *ago3* mutant alleles *ago3*^*t2*^ and *ago3*^*t3*^, which contain premature stop codons (Li et al, 2009a). No GSC loss was recorded in the mutant combination *ago3*^*t2*/*t3*^ in 7-, 14-or 21-day-old females, showing that GSC self-renewal was not affected in the *ago3* mutant. In contrast, *ago3*^*t2*/*t3*^ females showed a prominent defect in GSC differentiation, with a large proportion of germaria having a higher number of undifferentiated germ cells with a spectrosome (2 to 4 in control germaria, versus 6 to 9 in *ago3*^*t2*/*t3*^ germaria) (Supplementary Figure 2).

Together, these results demonstrate that Aub is required intrinsically in GSCs for their self-renewal and differentiation. Aub maintains GSCs by preventing their differentiation independently of Bam expression. In contrast, Ago3 is specifically involved in GSC differentiation.

### Aub function in GSC self-renewal is independent of Chk2

The Chk2-dependent DNA damage checkpoint is activated in several piRNA pathway mutants, leading to developmental defects during mid-oogenesis and, in turn, defective dorsoventral and anteroposterior embryonic patterning (Klattenhoff et al, 2007; Pane et al, 2007). These developmental defects are partially rescued in double mutants of Chk2 kinase (*mnk* mutant) and different piRNA pathway components. DNA damage in piRNA pathway mutants is thought to result from transposable element transposition. Mobilization of *P*-elements in crosses that induce hybrid dysgenesis (*i.e.* the crossing of females devoid of *P*-elements with males that contain *P*-elements) leads to a block in GSC differentiation, which is partly rescued by mutation in Chk2 (Rangan et al, 2011). To address whether the defects in GSC self-renewal and differentiation in *aub* mutant ovaries might be due to activation of the Chk2-dependent checkpoint, we analyzed *mnk*^*p6*^*aub*^*HN2/QC42*^ double mutant ovaries using anti-Hts and anti-Vasa immunostaining. The *aub* mutant tumor phenotype of undifferentiated cell accumulation was almost completely rescued by *mnk*^*p6*^, demonstrating that this phenotype depends on Chk2 (Figure 2A-C). Strikingly, the GSC loss phenotype was not robustly rescued by *mnk*^*p6*^, showing that this defect was mostly independent of Chk2 checkpoint activation (Figure 2A-C).

**Figure 2:**
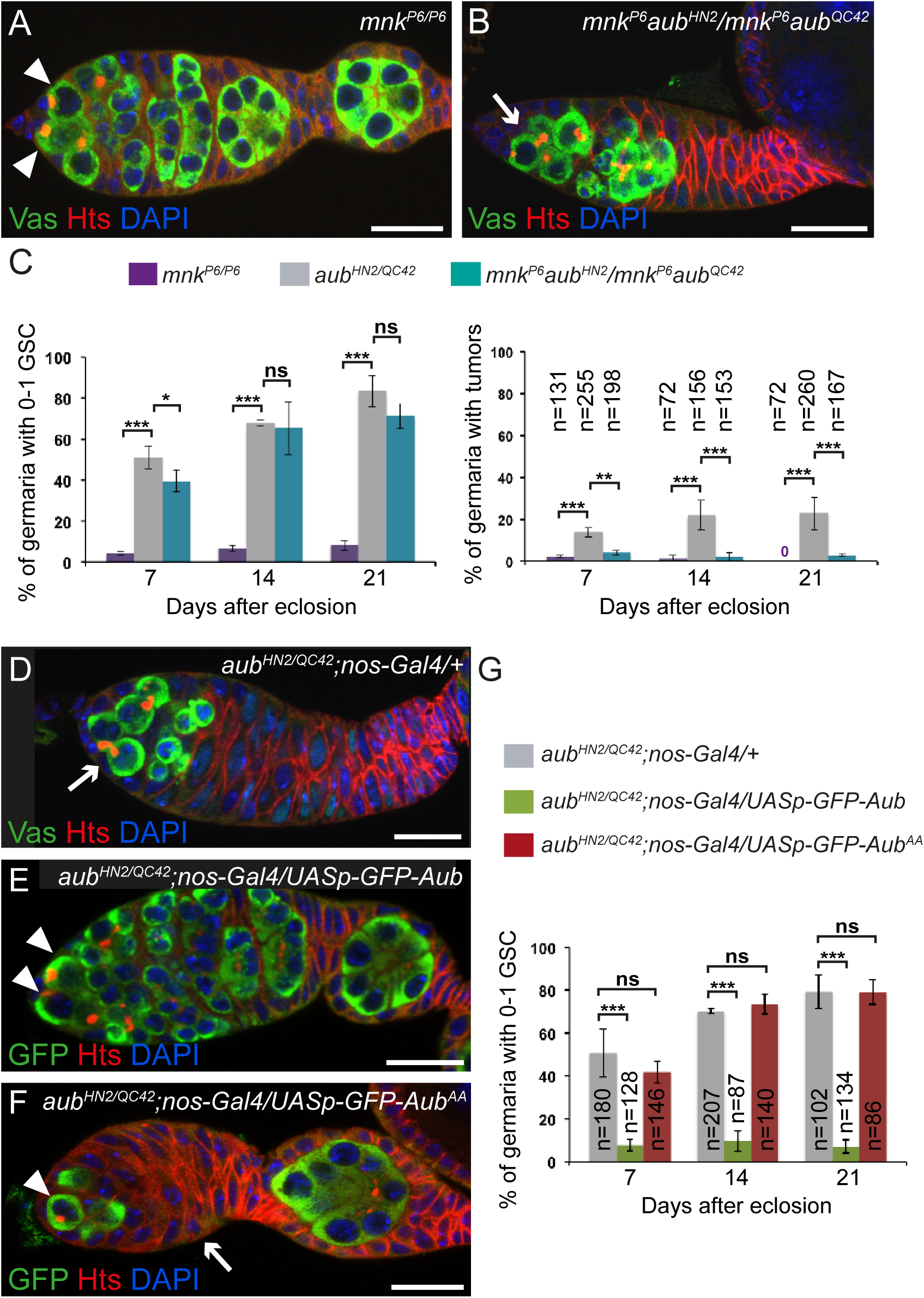
The role of Aub in GSC self-renewal requires its loading with piRNAs, but is independent of Chk2. (A-B) Immunostaining of germaria with anti-Vasa (green) and anti-Hts (red). DAPI (blue) was used to visualize DNA. Examples of *mnk*^*P6*^ and *mnk*^*P6*^*aub*^*HN2*^*/mnk*^*P6*^*aub*^*QC42*^ germaria are shown. White arrowheads indicate GSCs; the white arrow indicates GSC loss. (C) Quantification of mutant germaria with 0 to 1 GSC, or with GSC tumors in the indicated genotypes. The number of scored germaria (n) is indicated on the right graph. *** *p*-value <0.001, * *p*-value <0.05, ns: non significant, using the χ2 test. (D-F) Immunostaining of germaria from 7 day-old females with anti-Vasa (green) and anti-Hts (red), or with anti-GFP (green) and anti-Hts (red). DAPI (blue) was used to visualize DNA. *aub*^*HN2*^*/aub*^*QC42*^*; nos-Gal4/+* was used as a negative control. Examples of rescue in *aub*^*HN2*^*/aub*^*QC42*^*; nos-Gal4/UASp-GFP-Aub* germarium (E), and of lack of rescue in *aub*^*HN2*^*/aub*^*QC42*^*; nos-Gal4/UASp-GFP-Aub*^*AA*^ germarium (F). White arrowheads indicate GSCs; white arrows indicate GSC loss in (D) and germ cell loss in (F). (G) Quantification of mutant germaria with 0 to 1 GSC shown in (D-F). The number of scored germaria (n) is indicated. *** *p*-value <0.001, ns: non-significant, using the χ2 test. Scale bar: 10 μm in A-B and D-F.

These results show that the mild defect in GSC differentiation in the *aub* mutant is due to activation of the DNA damage checkpoint. In contrast, the prominent GSC self-renewal defect does not arise from activation of the DNA damage checkpoint, strongly supporting a more direct role of Aub in GSC self-renewal.

### Aub loading with piRNAs is required for GSC self-renewal

To determine whether the role of Aub in GSC self-renewal depends on its loading with piRNAs, we used an Aub double point mutant in the PAZ domain that is unable to bind piRNAs (Aub^AA^) (Barckmann et al, 2015). In contrast to *UASp-GFP-Aub,* which was able to rescue the GSC loss phenotype in *aub* mutant flies when expressed with *nos-Gal4* (Figure 1E, F), expression of *UASp-GFP-Aub*^*AA*^ at similar levels (Supplementary Figure 3A-C) failed to rescue this phenotype (Figure 2D-G). These data indicate that Aub loading with piRNAs is required for GSC self-renewal. To confirm this result we used a piRNA pathway mutant in which piRNA biogenesis is strongly compromised. *armitage* (*armi*) encodes a RNA helicase required for piRNA biogenesis (Cook et al, 2004; Malone et al, 2009). Immunostaining of *armi*^*1/72*^ ovaries with anti-Hts and anti-Vasa revealed a GSC loss that increased with time (Supplementary Figure 3D-F), showing a function for *armi* in GSC self-renewal.

We conclude that Aub function in GSC self-renewal depends on its loading with piRNAs.

### Aub interacts with the CCR4-NOT complex for GSC self-renewal

Previous reports have shown that PIWI proteins can recruit the CCR4-NOT deadenylation complex to repress mRNAs at the post-transcriptional level (Gou et al, 2014; Rouget et al, 2010). To address whether Aub might act through a similar mode of action in GSCs, we analyzed Aub and CCR4 colocalization. CCR4 is present diffusely in the cytoplasm and accumulates in cytoplasmic foci, in GSCs (Joly et al, 2013). Aub also has a diffuse distribution in the cytoplasm and is present in foci that surround the nucleus collectively referred to as “nuage” (Harris & Macdonald, 2001).

Colocalization occurred in diffusely distributed pools of proteins and occasionally in foci (Figure 3A-A’’), consistent with deadenylation not taking place in foci (Joly et al, 2013). We then overexpressed CCR4-HA in GSCs using *nos-Gal4* and found that CCR4-HA was able to recruit Aub in discrete cytoplasmic regions where CCR4-HA had accumulated, consistent with the presence of CCR4 and Aub in the same complex in GSCs (Figure 3B-B’’). Coimmunoprecipitation experiments in early ovaries from 1-day-old females revealed that GFP-Aub was able to coprecipitate the NOT1 and CCR4 subunits of the CCR4-NOT deadenylation complex, either in the presence or absence of RNA (Figure 3C).

**Figure 3:**
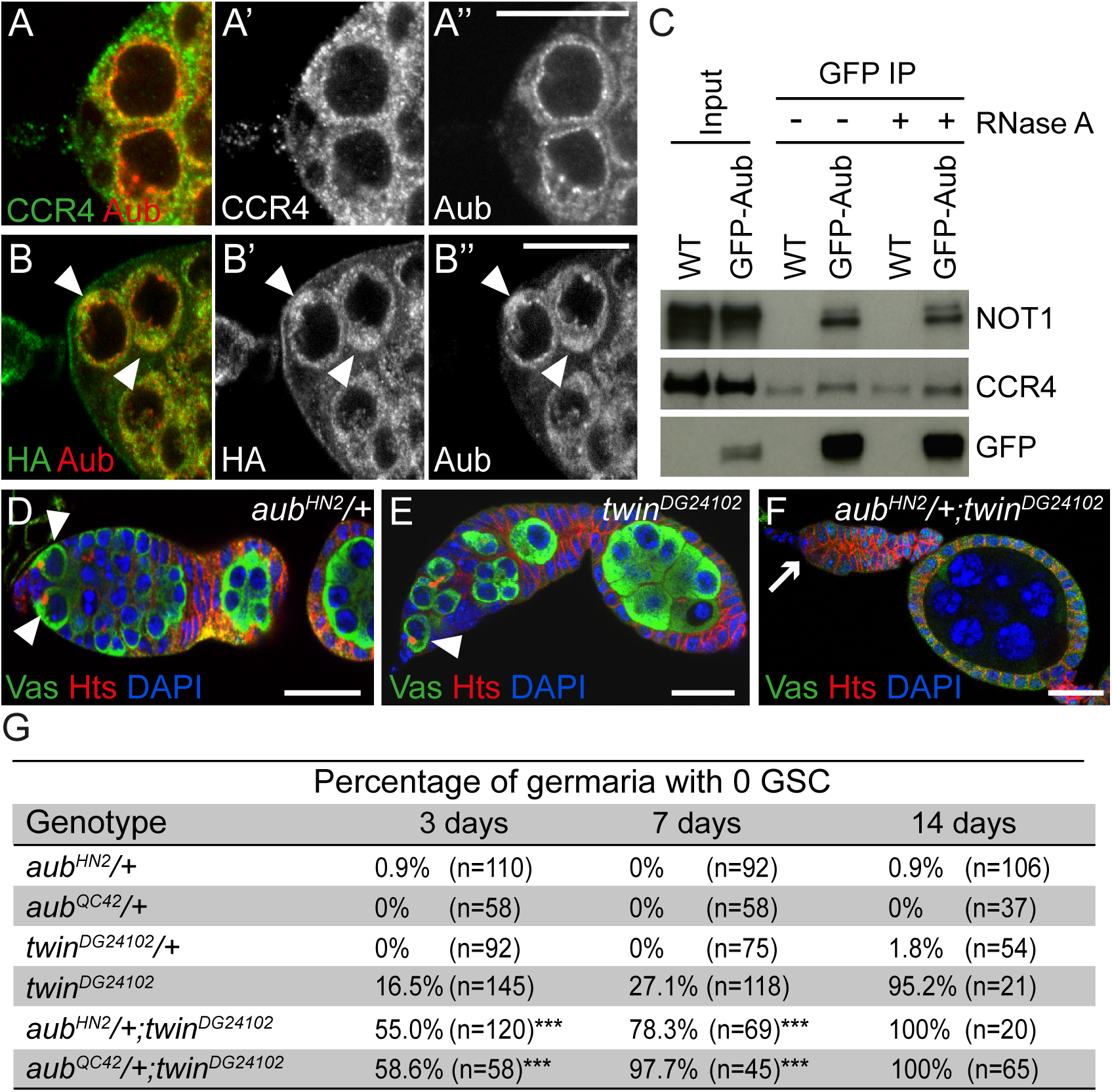
Physical and genetic interaction between Aub and the CCR4 deadenylase. (A-B’’) Immunostaining of wild-type (A-A’’) or *nos-Gal/UASp-CCR4-HA* (B-B’’) germaria with anti-Aub (red), and anti-CCR4 or anti-HA (green), respectively. GSCs are shown in (A-A’’). White arrowheads in (B-B’’) indicate cytoplasmic accumulation of CCR4-HA colocalized with Aub in GSCs. (C) Co-immunoprecipitation (IP) of CCR4 and NOT1 with GFP-Aub in ovaries. Wild-type (WT, mock IP) or *nos-Gal4/UASp-GFP-Aub* (GFP-Aub) ovarian extracts were immunoprecipitated with anti-GFP, either in the absence or the presence of RNase A. Western blots were revealed with anti-GFP, anti-CCR4 and anti-NOT1. Inputs are 1/10 of extracts prior to IP. (D-F) Genetic interaction between *aub* and *twin* in GSC self-renewal. Immunostaining of germaria with anti-Vasa (green) and anti-Hts (red). DAPI (blue) was used to visualize DNA. Examples of *aub*^*HN2/+*^, *twin*^*DG24102*^ and *aub*^*HN2/+*^; *twin*^*DG24102*^ germaria are shown. White arrowheads indicate GSCs; the white arrow indicates GSC loss. (G) Quantification of mutant germaria with no GSC in 3, 7 and 14 day-old females of the genotypes shown in (D-F). The number of scored germaria (n) is indicated. *** *p*-value <0.001 using the χ2 test. Scale bar: 10 μm in A-B and D-F.

We previously reported that the *twin* gene encoding the CCR4 deadenylase is essential for GSC self-renewal (Joly et al, 2013). To genetically determine if *aub* acts together with *twin* in GSC self-renewal, we tested whether GSC loss in the hypomorphic allele *twin*^*DG24102*^ might be enhanced by reducing the gene dosage of *aub*. GSC loss in *twin*^*DG24102*^ was accelerated in the presence of heterozygous *aub*^*HN2*^ or *aub*^*QC42*^ mutations, consistent with a role for Aub and CCR4 in the same molecular pathway for GSC self-renewal (Figure 3D-G).

Together, these results show that Aub and CCR4 are in complex in GSCs, and cooperate for GSC self-renewal.

### *Cbl* is an mRNA target of Aub in GSCs

To identify mRNA targets of Aub in GSCs, we looked for candidate genes with a reported role in GSC biology or other stem cell lineages (Supplementary Figure 4A). Eight genes were selected, five of which produce mRNAs that directly interact with Aub in embryos (Barckmann et al, 2015).

Antibody staining in ovaries containing clonal *aub* mutant GSCs was used to record potential increased levels of the corresponding proteins in mutant GSCs as compared to control (Supplementary Figure 4A-C’’). We found a mild increase in Mei-P26 and Fused protein levels in *aub* mutant GSCs, and a more prominent increase in Nos levels (Supplementary Figure 4A-C’’). Increased Nos protein levels in *aub* mutant GSCs suggests that the direct regulation of *nos* mRNA by Aub occurring in the early embryo is maintained in other biological contexts (Rouget et al, 2010).

Cbl protein displayed the highest increased levels in *aub* mutant GSCs compared to control. We thus focused on the possible regulation of the *Cbl* proto-oncogene by Aub. *Cbl* encodes two isoforms through alternative splicing: a long isoform (CblL) and a short isoform (CblS), both of which contain the N-terminal phosphotyrosine binding domain that binds phosphotyrosine kinases, in addition to a ring finger domain that acts as an E3 ubiquitin ligase (Figure 4A) (Robertson et al, 2000). We used two available monoclonal antibodies directed against either the long Cbl isoform (8C4) or both isoforms (10F1) (Pai et al, 2006), to analyze the deregulation of *Cbl* in *aub* mutant GSCs. Cbl protein levels were significantly increased in *aub* mutant GSCs as observed with either antibody, although a stronger effect was revealed with the 8C4 antibody (specific to CblL) (Figure 4B-E). These results suggest the regulation of *CblL* mRNA by Aub in the GSCs and are consistent with the reported mRNA expression of *CblL* in germaria and *CblS* in later stages of oogenesis (Pai et al, 2006).

**Figure 4:**
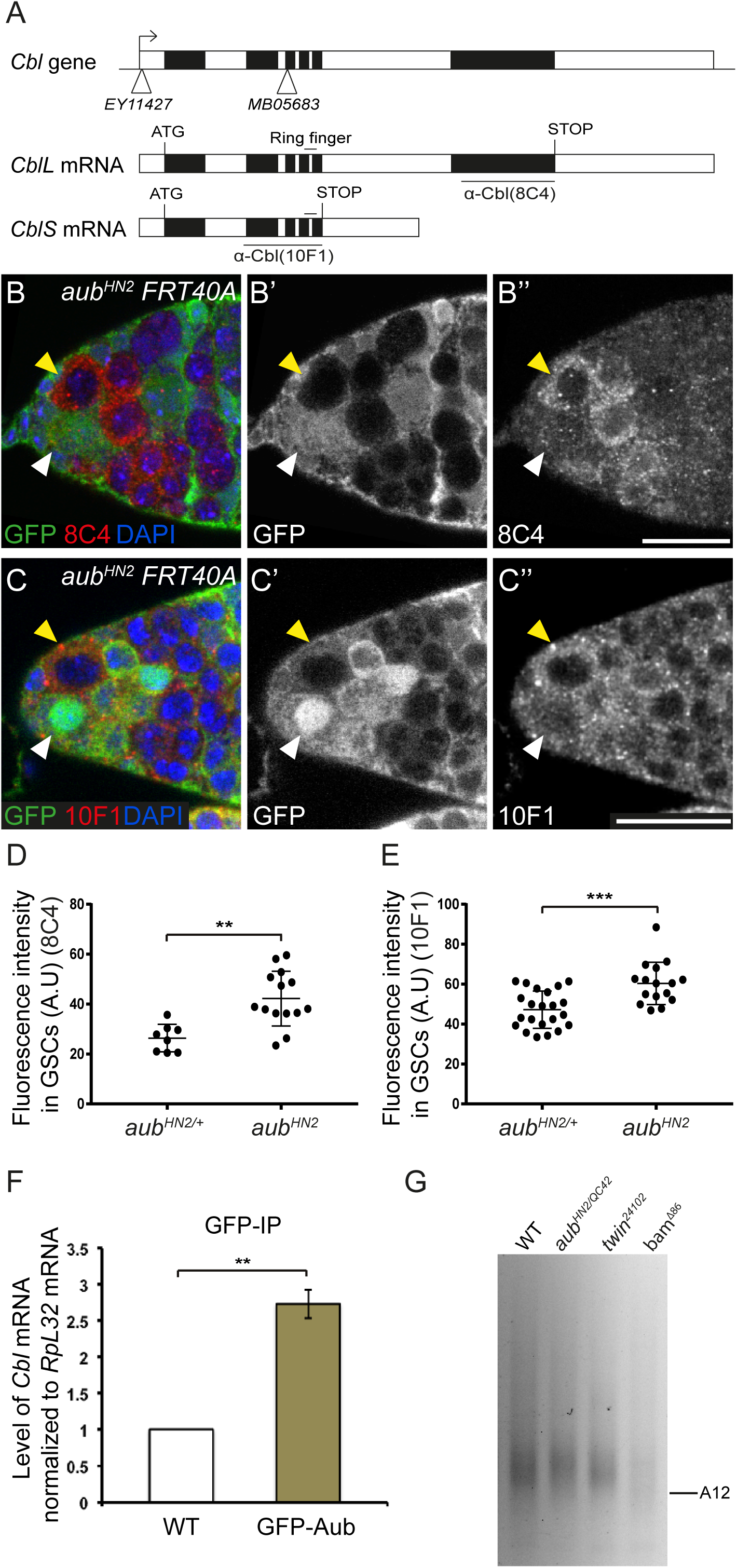
Aub represses *Cbl* expression in the GSCs. (A) Genomic organization of the *Cbl* locus and *Cbl* mRNAs. Open boxes represent UTRs and introns, black boxes are exons. The insertion points in the *Cbl*^*EY11427*^ and *Cbl*^*MB05683*^ mutants are represented by white triangles. The region encoding the E3 ubiquitin ligase domain (Ring finger) is indicated. The regions used to raised the 8C4 and 10F1 monoclonal antibodies are underlined. (B-C’’) Immunostaining of mosaic germaria with anti-GFP (green), to identify clonal cells by the lack of GFP, and either 8C4 (B-B’’) or 10F1 (C-C’’) monoclonal anti-Cbl (red). DAPI was used to visualize DNA. White arrowheads indicate *aub*^*HN2/+*^ control GSCs; yellow arrowheads indicate clonal mutant *aub*^*HN2*^ GSCs. (D-E) Quantification of Cbl protein levels in *aub*^*HN2/+*^ and *aub*^*HN2*^ mutant GSCs using fluorescence intensity of immunostaining with 8C4 or 10F1. Fluorescence intensity was measured in arbitrary units using the ImageJ software. Horizontal bars correspond to the mean and standard deviation. \*\*\**p*-value <0.001 using the Student’s *t* test. (F) RNA immunoprecitation (IP) with anti-GFP antibody in wild-type (mock IP) and *nos-Gal4/UASp-GFP-Aub* ovarian extracts. *Cbl* mRNA was quantified using RT-qPCR. Normalization was with *RpL32* mRNA. Mean of three biological replicates. The error bar represents standard deviation. \*\**p*-value <0.01 using the Student’s *t* test. (G) ePAT assay of *CblL* mRNA. Ovaries from 1-day-old (germarium to stage 8) wild-type, *aub* and *twin* mutant females, and from 4-to 7-day-old *bam* mutant females were used.

We used RNA immunoprecipitation with Aub to confirm the potential regulation of *Cbl* by Aub at the mRNA level. GFP-Aub protein was immunoprecipitated from *UASp-GFP-Aub/nos-Gal4* or wild-type (mock immunoprecipitation) ovaries. Quantification of *Cbl* mRNA by RT-qPCR revealed that it is enriched in GFP-Aub immunoprecipitates as compared to the mock immunoprecipitates (Figure 4F). Consistent with Aub interaction with *Cbl* mRNA, Aub iCLIP experiments in 0-2 embryos have revealed the direct binding of Aub to *Cbl* mRNA (Barckmann et al, 2015). To address the role of piRNAs in the regulation of *Cbl* mRNA by Aub, we analyzed Cbl protein levels in GSCs from *armi* mutant females, in which piRNA biogenesis is affected. Immunostaining of *armi* mutant germaria with anti-Cbl antibody 8C4 revealed a significant increase of CblL levels in *armi* mutant GSCs (Supplementary Figure 3G-I).

We then analyzed the potential deregulation of *Cbl* mRNA in the absence of CCR4 deadenylase. We performed Cbl immunostaining in ovaries containing clonal *twin* mutant GSCs using the 8C4 antibody. Similar to the results observed in *aub* mutant GSCs, the levels of CblL isoform were increased in *twin* mutant GSCs (Supplementary Figure 5). To address whether the regulation of *Cbl* mRNA by Aub and CCR4 occurred at the level of poly(A) tail length, we measured the poly(A) tail of *CblL* mRNA in ovaries using ePAT assays. ePAT assays from *bam*^*δ86*^ ovaries that only contained undifferentiated GSC-like cells confirmed the presence of the long *CblL* mRNA in these cells (Figure 4G). *CblL* poly(A) tails were not notably affected in *twin* and *aub* mutant early ovaries as compared to the wild-type, suggesting that *CblL* mRNA regulation by Aub/CCR4-NOT does not involve deadenylation. Indeed, the CCR4-NOT complex has the capacity to repress mRNA translation independently of its role in deadenylation, through the recruitment of translational repressors (Chekulaeva et al, 2011; Chen et al, 2014; Mathys et al, 2014).

Taken together, these results reveal *Cbl* mRNA as a direct target of Aub/CCR4-NOT-dependent translational repression, and the implication of piRNAs in this regulation.

### Regulation of *Cbl* mRNA by Aub is required for GSC self-renewal

To determine whether translational repression of *Cbl* mRNA by Aub is relevant to GSC self-renewal, we analyzed the function of *Cbl* in GSCs. We first determined the expression pattern of Cbl in the germarium using immunostaining. The *bamP-GFP* reporter was used to mark the cystoblast to 8-cell cysts (Chen & McKearin, 2003). Co-staining of both Cbl antibodies with GFP revealed the presence of Cbl at low levels in GSCs, cystoblasts and early (2-cell) cysts. This was followed by reduced levels in the remaining dividing and differentiating cysts up to region 2a of the germarium, and stronger expression starting in region 2b in both somatic and germ cells (Figure 5A-B’’). Strikingly, co-staining with anti-Cbl and anti-GFP antibodies of germaria expressing GFP-Par1 to visualize the spectrosomes, revealed a significant increase, albeit slight and transient, of Cbl levels in cystoblasts as compared to GSCs (Figure 5C, D, Supplementary Figure 6).

**Figure 5:**
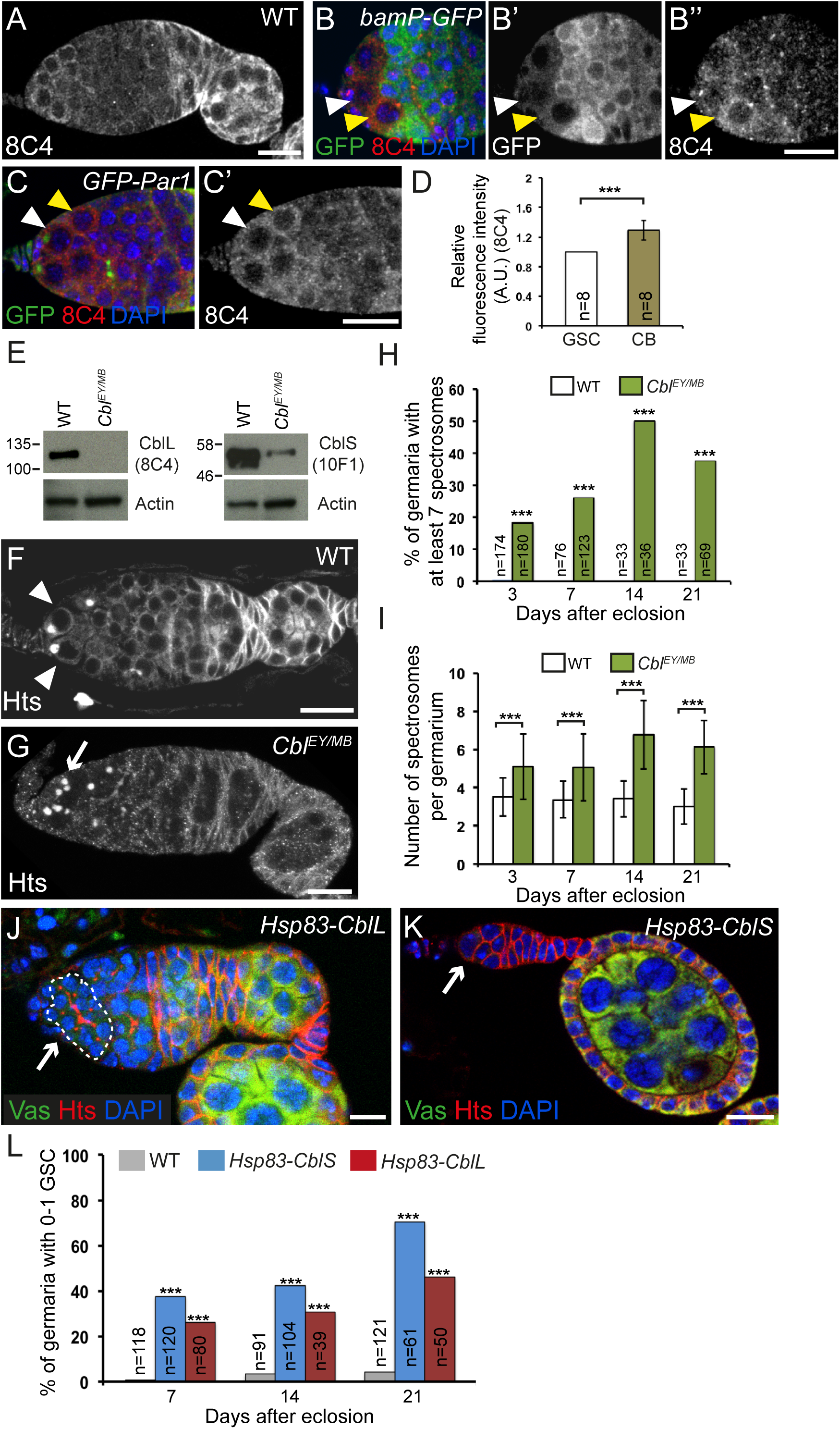
*Cbl* is required for GSC differentiation. (A-D) *Cbl* expression in germaria. (A) Immunostaining of wild-type germaria with anti-Cbl 8C4. (B-B’’) Immunostaining of *bamP-GFP* germaria that express GFP under the *bam* promoter, with anti-GFP (green) and anti-Cbl 8C4 (red). (C-C’) Immunostaining of germaria expressing GFP-Par1 to label spectrosomes with anti-GFP (green) and anti-Cbl 8C4 (red). DAPI (blue) was used to visualize DNA. White arrowheads indicate GSCs; yellow arrowheads indicate cystoblasts. (D) Quantification of Cbl protein levels in GSCs and cystoblasts using fluorescence intensity of immunostaining with 8C4 antibody. Fluorescence intensity was measured in arbitrary units using ImageJ. The number of scored cells (n) is indicated. Intensity in GSCs was set to 1. The error bar indicates standard deviation. \*\*\**p*-value <0.001 using the Student’s *t* test. (E) Western blots of protein extracts from wild-type and *Cbl* mutant ovaries showing CblL (left panel) and CblS (right panel) revealed with 8C4 and 10F1 antibodies, respectively. A low level of CblS remained, while no CblL was present in the *Cbl*^*EY11427/MB05683*^ combination. The 10F1 antibody recognizes very poorly CblL in western blot (not shown). (F-I) *Cbl* is required for germline differentiation. Immunostaining of wild-type and *Cbl*^*EY11427/MB05683*^ germaria with anti-Hts to visualize spectrosomes and fusomes. White arrowheads indicate GSCs; the white arrow indicates increased number of spectrosomes. (H) Quantification of germaria with increased number of spectrosomes, and (I) quantification of spectrosomes per germarium. The number of scored germaria (n) is indicated in (H). \*\*\**p*-value <0.001 using the χ2 test in (H), and the Student’s *t* test in (I). (J-L) *Cbl* induces GSC differentiation. Immunostaining of germaria overexpressing *Cbl* with *Hsp83-CblL* (J) or *Hsp83-CblS* (K) with anti-Vasa (green) and anti-Hts (red). DNA (blue) was revealed with DAPI. White arrows indicate a cyst in the GSC niche (J, outlined) or the loss of GSCs and germ cells (K). (L) Quantification of mutant germaria with 0 to 1 GSC in the indicated genotypes. The number of scored germaria (n) is indicated. *** *p*-value <0.001 using the χ2 test.

Null alleles of *Cbl* are larval/pupal lethal (Pai et al, 2000; Pai et al, 2006). To address a potential role for *Cbl* in GSC biology in adults, we used two *Cbl* insertion alleles: *Cbl*^*EY11427*^, which contains a *P-UAS* insertion in the *Cbl* 5’UTR (Bellen et al, 2004), and *Cbl*^*MB05683*^, which contains a *Minos*-based insertion in the third exon (Metaxakis et al, 2005) (Figure 4A). *Cbl*^*EY11427/MB05683*^ transheterozygotes mostly died at the pupal stage, with a small number of adult escapers surviving for 3-4 days at 25°C. The number of escapers and their survival time increased at 22°C. Western blots of ovaries from these escapers revealed that the CblL isoform was absent in this mutant combination, and that CblS levels were strongly reduced (Figure 5E). Anti-Hts immunostaining of ovaries from 3-, 7-, 14-and 21-day-old *Cbl*^*EY11427*/*MB05683*^ mutant females revealed a defect in GSC differentiation. The number of undifferentiated germ cells containing a spectrosome in *Cbl* mutant germaria was higher than in the wild-type and increased with time (2 to 4 spectrosomes in the wild-type versus 6 to 9 in *Cbl* mutant) (Figure 5F-I).

In a reverse experiment, we overexpressed the long or short isoforms of Cbl in germ cells using the *Hsp83-CblL* and *Hsp83-CblS* transgenes (Pai et al, 2006) and analyzed germaria using immunostaining with anti-Hts and anti-Vasa. Consistent with a role for *Cbl* in GSC differentiation, overexpressing *Cbl* in the germline led to a GSC loss that increased with time (Figure 5J-L). These results identify a new *Cbl* function in GSC homeostasis.

Finally, we addressed whether the translational repression of *Cbl* mRNA by Aub in the GSCs has a functional role in their self-renewal. If increased Cbl protein levels in *aub* mutant ovaries contribute to the *aub* GSC loss phenotype, we would expect to reduce this phenotype by reducing *Cbl* gene dosage. We used both *Cbl*^*EY11427*^ and *Cbl*^*MB05683*^ heterozygous mutants in combination with *aub*^*HN2/QC42*^ and examined the germaria of these females at three time points by immunostaining with anti-Hts and anti-Vasa. Strikingly, the GSC loss phenotype in *aub* mutants was significantly rescued in the presence of both *Cbl* heterozygous mutants (Figure 6A-C). These results show that the regulation of *Cbl* by Aub is essential for GSC self-renewal.

**Figure 6:**
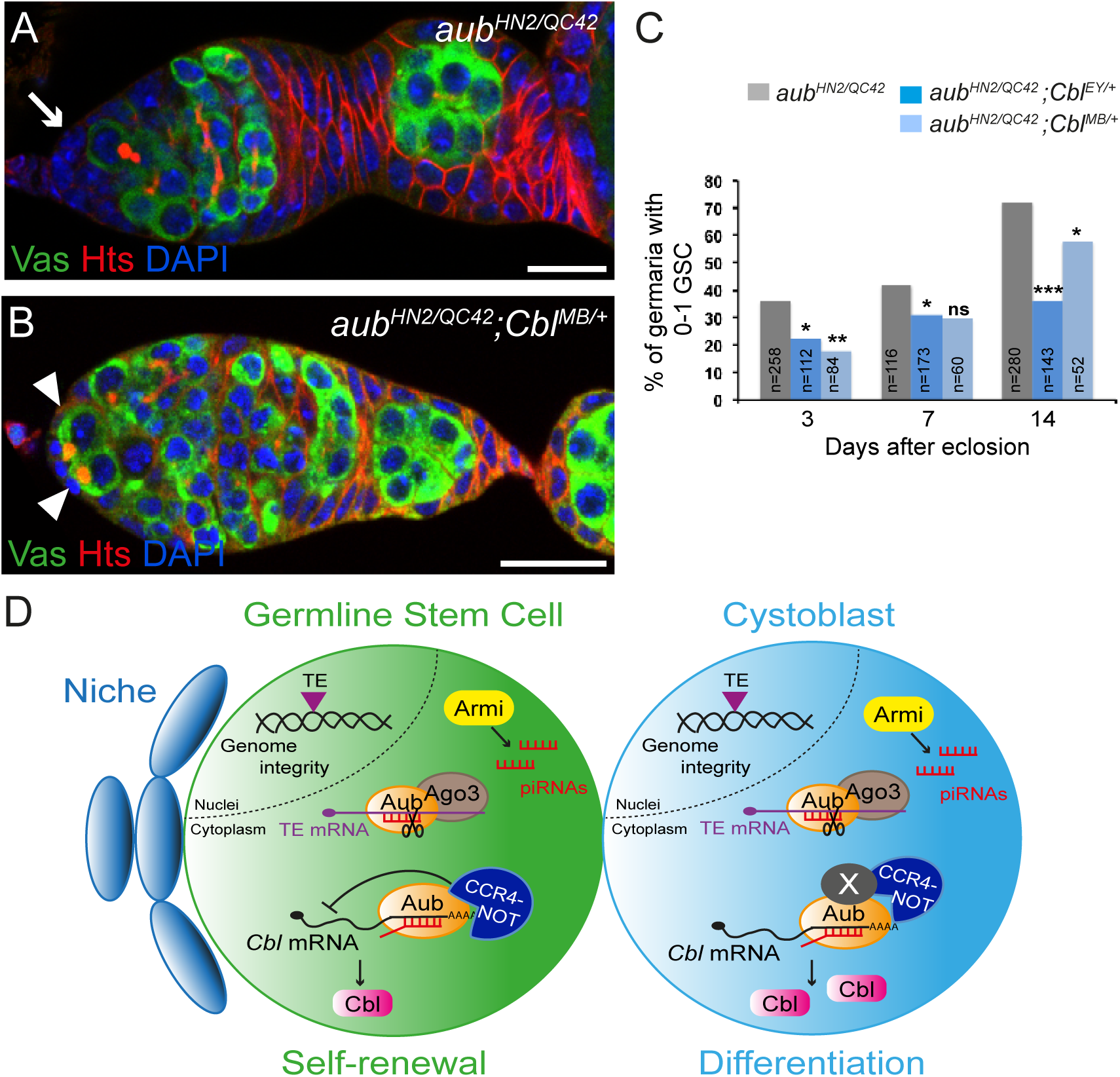
Regulation of Cbl by Aub in the GSCs is essential for their self-renewal. (A-B) Immunostaining of germaria from *aub*^*HN2/QC42*^ (A) and *aub*^*HN2/QC42*^*; Cbl*^*MB/+*^ (B) females with anti-Vasa (green) and anti-Hts (red). DAPI (blue) was used to visualize DNA. The white arrow indicates the lack of GSCs; white arrowheads indicate GSCs. (C) Quantification of mutant germaria with 0 to 1 GSC in the indicated genotypes. The number of scored germaria (n) is indicated. *** *p*-value <0.001, ** *p*-value <0.01, \**p*-value <0.05, ns: non-significant using the χ2 test. (D) Model of Aub function in GSCs. Aub is required intrinsically in GSCs for their self-renewal and differentiation. Aub function in self-renewal depends on translational repression of *Cbl* mRNA in GSCs through the recruitment of the CCR4-NOT complex. In cystoblasts, this translational repression is decreased, likely through the implication of at least another factor (X). As is the case for other translational controls in the GSC lineage, Aub/CCR4-NOT acts in fine-tuning Cbl levels. Aub function in GSC differentiation depends on activation of the Chk2 DNA damage checkpoint, consistent with a role in transposable element (TE) repression to maintain genome integrity; Ago3 has the same role in GSC differentiation.

Together, these data identify a new role for *Cbl* in GSC differentiation and reveal an essential role of *Cbl* mRNA translational repression by Aub for GSC self-renewal.

## Discussion

Here we have demonstrated that *aub* is required intrinsically in GSCs for their self-renewal. Our results show that this *aub* function does not depend on activation of the DNA damage response mediated by Chk2 kinase, but rather on regulation of protein-coding genes at the mRNA level. The main phenotype of *aub* mutant GSCs is a reduced capacity to self-renew, leading to their progressive loss by differentiation and this phenotype is not rescued by a mutant of the Chk2 kinase.

We provide evidence that the Aub mechanism of action involves the recruitment of the CCR4-NOT deadenylation complex (Figure 6D). Aub and the subunits of CCR4-NOT, NOT1 and CCR4 form a complex in the ovaries. Moreover, *aub* heterozygous mutants significantly increase the GSC loss phenotype of a *twin* hypomorphic mutant. Translational repression plays an essential role in GSC biology, for both GSC self-renewal and differentiation. Through their interaction with CCR4-NOT, the translational repressors Nos and Pum repress the translation of differentiation factors in GSCs for their self-renewal (Joly et al, 2013); in turn, Pum and Brat recruit CCR4-NOT in the cystoblasts for their differentiation by repressing the translation of self-renewal factors (Harris et al, 2011; Newton et al, 2015). Therefore, the CCR4-NOT complex is central to mRNA regulation for GSC homeostasis. We have identified Aub as a novel interactor of the CCR4-NOT deadenylation complex involved in translational repression of *Cbl* mRNA for GSC self-renewal. Interestingly, repression of *Cbl* mRNA by Aub/CCR4-NOT does not involve poly(A) tail shortening. This is consistent with the reported role of the CCR4-NOT complex in translational repression, independent of deadenylation. In this mode of regulation, the CCR4-NOT complex serves as a platform to recruit translational repressors such as DDX6/Me31B and Cup (Chen et al, 2014; Igreja & Izaurralde, 2011; Mathys et al, 2014). In addition, a role in translational regulation has been proposed for two mouse PIWI proteins, MILI and MIWI. MILI associates with the translation factor eIF3A, while both MILI and MIWI associate with the cap-binding complex (Grivna et al, 2006; Unhavaithaya et al, 2009). Similarly, Aub might also regulate mRNA translation through direct interaction with translation factors.

A major point addressed here is the characterization of the role of the piRNA pathway in GSC biology; until now, only the function of Piwi has been thoroughly analyzed in GSCs. Piwi is involved in somatic niche cells for GSC differentiation, as well as in GSCs for their maintenance and differentiation (Cox et al, 1998; Cox et al, 2000; Jin et al, 2013; Ma et al, 2014). Strikingly, the molecular mechanisms underlying the somatic role of Piwi for GSC differentiation are not related to transposable element repression, but gene regulation. Piwi interacts with components of the PRC2 complex, thus limiting PRC2 binding to chromatin and transcriptional repression (Peng et al, 2016). Piwi also represses *c-Fos* mRNA via its processing into piRNAs (Klein et al, 2016). Cutoff, a component of the piRNA pathway required for piRNA production, is reported to play a role in GSC self-renewal and differentiation, with a partial rescue of the differentiation defects by mutation in the Chk2 kinase (Chen et al, 2007; Pane et al, 2011). Here, we describe the GSC phenotypes in three additional piRNA pathway mutants: *aub*, *ago3*, and *armi* (Figure 6D). Notably, the three mutants display different defects in GSC biology. Specifically, *ago3* mutants only have differentiation defects, whereas the most prominent phenotype of *aub* and *armi* mutants is GSC loss. This suggests that different molecular pathways affect GSC biology in these mutants. Importantly, the effect of transposition *per se* on GSC homeostasis has been analyzed using *P*-element mobilization in PM hybrid dysgenesis crosses, and has been shown to induce defects in GSC differentiation (Rangan et al, 2011). These defects are partly rescued by Chk2 mutation, indicating that they arise following DNA damage. Accordingly, differentiation defects in *aub* mutant GSCs are also largely rescued by Chk2 mutation, and might result from transposition. Remarkably, GSC loss in the *aub* mutant was not rescued by the *Chk2* mutant. These results support the conclusion that defects in GSC self-renewal, the main phenotype in *aub* mutants, do not predominantly depend on transposable element mobilization or DNA damage, and are consistent with our identification of the *aub*-dependent regulation of a cellular mRNA for GSC self-renewal. These findings broaden the developmental functions of Aub and piRNAs as regulators of gene expression in various biological processes, and highlight their key role in developmental transitions.

Importantly, our study reveals a new function for Cbl in GSC biology. Overexpression and mutant analysis show the implication of *Cbl* in GSC differentiation. *Cbl* is a tumor suppressor gene encoding an E3 ubiquitin ligase that binds and represses receptor tyrosine kinases. In particular, Cbl regulates epidermal growth factor receptor (Egfr) signaling through ubiquitination and degradation of activated Egfr (Mohapatra et al, 2013). In *Drosophila*, the regulation of Gurken/Egfr signaling by Cbl is involved in dorsoventral patterning during oogenesis (Chang et al, 2008; Pai et al, 2000; Pai et al, 2006). Mutations in *Cbl* that disrupt E3 ubiquitin ligase activity lead to myeloid neoplasms in humans (Sanada et al, 2009). Indeed, Cbl plays a key role in hematopoietic stem cell homeostasis, in the maintenance of their quiescence and their long-term self-renewal capacity (An et al, 2015). Our data thus add a biological function for Cbl to yet another stem cell lineage.

Other E3 ubiquitin ligases are known to play important roles in GSC biology. Mei-P26 and Brat, two members of the conserved Trim-NHL family of proteins, contain E3 ubiquitin ligase domains and have roles in stem cell lineages. Mei-P26 in particular is involved in GSC self-renewal and differentiation, and this dual function partly depends on a very tight regulation of its levels in these cells (Joly et al, 2013; Li et al, 2012; Neumuller et al, 2008). Smurf is another E3 ubiquitin ligase that plays a major role in GSC differentiation. Specifically, Smurf associates with the Fused serine/threonine kinase in cystoblasts to degrade the Thickveins receptor and thus repress BMP signaling, the main signaling pathway in the GSC lineage. This mechanism generates a steep gradient of BMP activity between GSCs and cystoblasts (Xia et al, 2010). Cbl might participate in the regulation of Egfr signaling or other signaling pathways in the GSC lineage. Although the role of Egfr in adult GSC biology in the ovary has not yet been addressed, this signaling pathway is involved in regulating the primordial germ cell number in the larval gonad (Gilboa & Lehmann, 2006) and in the GSC mitotic activity in adult males (Parrott et al, 2012).

Our data highlight an important role of Aub in fine-tuning Cbl levels for GSC self-renewal. Translational regulation is known to be central for cell fate choices in adult stem cell lineages. RNA binding proteins and microRNAs have a recognized regulatory function in female GSCs to trigger cell fate changes through cell-specific regulation of mRNA targets. Here, we reveal piRNAs as an additional layer of translational regulators for GSC biology. PIWI proteins and piRNAs are stem cell markers in somatic stem cells of higher organisms, as well as in pluripotent stem cells involved in regeneration in lower organisms (Juliano et al, 2011). This function of piRNAs in translational control is likely to be conserved in these stem cell lineages and might play a key role in stem cell homeostasis.

PIWI proteins and piRNAs are upregulated in a number of cancers, and functional studies in *Drosophila* have shown that this upregulation participates in cancer progression (Fagegaltier et al, 2016; Janic et al, 2010). This suggests that the role of piRNAs in the translational control of cellular mRNA targets might also be crucial to cancer.

## Materials and Methods

### *Drosophila* stocks and genetics

*Drosophila* stocks are described in Expanded View Materials and Methods.

## Immunostaining and image analysis

Ovaries were dissected at room temperature in PBS supplemented with 0.1% Tween-20 (PBT), fixed with 4% paraformaldehyde, blocked with PBS containing 10% BSA for 1 h and incubated in primary antibodies with PBT 1% BSA overnight at 4°C. Primary antibodies were then washed three times with PBT 1% BSA for 10 min at room temperature. Secondary antibodies were diluted in PBT 0.1% BSA and were incubated for 4 h at room temperature. Secondary antibodies were then washed three times in PBT for 10 min. Primary antibodies were used at the following concentrations: mouse anti-Hts (1B1; Developmental Studies Hybridoma Bank (DSHB), University of Iowa) 1/100; rabbit anti-Vasa (Santa Cruz Biotechnology) 1/1000; rat anti-Vasa (DSHB) 1/50; rabbit anti-cleaved Caspase 3 (Biolabs) 1/300; rabbit anti-Bam (a gift from D. Chen) 1/2000; rabbit anti-Aub (ab17724; Abcam) 1/500; mouse anti-Aub (4D10; (Gunawardane et al, 2007)) 1/1500; mouse anti-HA (ascite produced from 12CA5 (Joly et al, 2013)) 1/2000; rabbit anti-GFP (A6455; Invitrogen) 1/500; mouse anti-Cbl (8C4; DSHB) 1/100; mouse anti-Cbl (10F1; a gift from LM. Pai) 1/300; rabbit anti-Nanos (a gift from A. Nakamura), 1/1000; rabbit anti-Brat (Betschinger et al, 2006) 1/300; rabbit anti-Mei-P26 (Liu et al, 2009) 1/100; mouse anti-Fused (22F10; DSHB) 1/100; rabbit anti-Pgc (Hanyu-Nakamura et al, 2008) 1/1000; rabbit anti-Bruno (Sugimura & Lilly, 2006) 1/3000; rabbit anti-Lola (a gift from E. Giniger (Giniger et al, 1994)) 1/100. Secondary antibodies (Alexa 488-and Cy3-conjugated; Jackson ImmunoResearch) were used at 1/300. DNA staining was performed using DAPI at 0.5 μg/mL. Images were captured with a Leica SP8 Confocal microscope and analyzed using the ImageJ software. Fluorescence intensity was measured with ImageJ software in wild-type, heterozygous or mutant GSCs. For each GSC, the mean fluorescence intensity was determined using three independent quantifications in three different cytoplasmic regions in the same confocal section; the number of cells analyzed (n) is indicated in the bar graphs for each genotype.

## Coimmunoprecipitations and western blots

Protein coimmunoprecipitations were performed using 60 ovaries per experiment from *w*^*1118*^ and *nos-Gal4/UASp-GFP-Aub* 3-day-old females. Ovaries were homogenized in 600 μL of DXB-150 (25 mM Hepes-KOH pH 6.8, 250 mM sucrose, 1 mM MgCl2, 1 mM DTT, 150 mM NaCl, 0.1% Triton X-100) containing cOmplete™ EDTA-free Protease Inhibitor Cocktail (Roche) and either RNase Inhibitor (0.25 U/μL; Promega) or RNase A (0.1 U/μL; Sigma). 50 μL of Dynabeads Protein G (Invitrogen) were incubated with 15 μL of mouse anti-GFP antibody (3E6; Invitrogen) for 1 h on a wheel at room temperature. Protein extracts were cleared on 30 μL of Dynabeads Protein G previously equilibrated with DXB-150 for 30 min at 4°C. The pre-cleared protein extracts were incubated with Dynabeads Protein G bound to mouse anti-GFP antibody for 3 h at 4°C. The beads were then washed 7 times with DXB-150 for 10 min at room temperature. Proteins were eluted in NUPAGE buffer supplemented with 100 mM DTT at 70°C. Western blots were performed as previously reported (Benoit et al, 1999) with antibodies used at the following concentrations: rabbit anti-GFP (Invitrogen) 1/1000; rabbit anti-CCR4 (Temme et al, 2004) 1/1000; mouse anti-NOT1 (Temme et al, 2010) 1/250; mouse anti-Cbl (8C4; DSHB) 1/1000; mouse anti-Cbl (10F1) 1/1000; rabbit anti-Actin (Sigma) 1/2500.

## RNA-immunoprecipitations and RT-qPCR

For RNA-immunoprecipitations, protein extracts were performed using 300 ovaries from *w*^*1118*^ and *nos-Gal4/UASp-GFP-Aub* females. Ovaries were homogenized in 600 μL of DXB-150 containing cOmplete™ EDTA-free Protease Inhibitor Cocktail (Roche) and RNase Inhibitor (0.25 U/μL; Promega). 50 μL of Protein G Mag Sepharose (GE Healthcare) were incubated with 2 μg mouse anti-GFP antibody (3E6; Invitrogen) for 3 h on a wheel at 4°C. Protein extracts were cleared on 50 μL of Protein G Mag Sepharose previously equilibrated with DXB-150 for 30 min at 4°C. The pre-cleared protein extracts were incubated with Protein G Mag Sepharose bound to mouse anti-GFP antibody for 3 h at 4°C. The beads were then washed 7 times with DXB-150 for 10 min at room temperature. RNA was prepared using TRIzol (Invitrogen), followed by DNA removal with TURBO DNA-free (Ambion). The total RNA amount was used for reverse transcription; RT-qPCR was performed with the LightCycler System (Roche Molecular Biochemical) using three independent RNA extractions and the primers listed in Expanded View Materials and Methods.

## Poly(A) tail assays

ePAT assays were performed as previously described (Chartier et al, 2015) using the primer indicated in Expanded View Materials and Methods.

## Author contributions

PRR performed the experiments and analyzed the data; AC performed PAT assays and coimmunoprecipitations; MS analyzed the data; PRR and MS designed the study and wrote the manuscript; all authors discussed the manuscript.

## Acknowledgements

We are very grateful to D Chen, E Giniger, A Gonzalez-Reyes, J Knoblich, A Nakamura, LM Pai, M Siomi, P Zamore for their gifts of fly stocks or antibodies. We thank AL Finoux and W Joly for initial work in this study. This work was supported by the CNRS UMR9002, ANR (ANR-2010-BLAN-1201 01 and ANR-15-CE12-0019-01), FRM (“Equipe FRM 2013 DEQ20130326534”). PRR held a salary from the Labex EpiGenMed/University of Montpellier and from the Fondation ARC. The authors declare that they have no conflict of interest.

